# Establishment and Molecular Characterization of Patient-Derived Organoids for Primary Central Nervous System Lymphoma

**DOI:** 10.1101/2024.10.21.619549

**Authors:** Shengjie Li, Jun Ren, Jianing Wu, Zuguang Xia, Yingzhu Li, Chengxun Li, Wenjun Cao

## Abstract

Primary central nervous system lymphoma (PCNSL) exhibits substantial heterogeneity, both intra-tumoral and intertumoral, posing challenges in developing effective treatment methods. Existing in vitro models fail to simulate the inherent microenvironment and the cellular and mutational diversity of native tumors and require a prolonged generation time. To address this concern, we described an organoid culture method for patient-derived PCNSL organoids (CLOs) and evaluated them through extensive molecular characterization. These CLOs accurately mimicked the histological attributes, gene expression landscapes and mutational profiles of their original tumors through rigorous histopathological analysis, RNA sequencing and whole-exome sequencing. Notably, CLOs were generated within 2 weeks, demonstrating rapid development and reliability. Furthermore, therapeutic profiling was performed on three selected CLOs using four standard treatment drugs. High concordance was observed between the drug responses of patients and those observed in the CLOs, with specific sensitivity to ibrutinib and methotrexate and resistance to dexamethasone and rituximab. Taken together, these results emphasize that CLOs can effectively emulate the key characteristics of PCNSL, enhancing the understanding of the genetic landscape of this complex disease. CLOs provide a rapid and reliable platform for exploring individualized treatment strategies, potentially accelerating the transition of research findings to clinical practice.

## Introduction

Primary central nervous system lymphoma (PCNSL) is an aggressive type of malignant brain tumour that is characterised as diffuse large B-cell lymphoma (DLBCL) in 95% of cases and has persistently high mortality rates [1,2]. Over the past two decades, the prognosis of PCNSL has shown significant improvement primarily owing to the widespread use of high-dose methotrexate (MTX) chemotherapy and consolidation treatment approaches including radiation therapy, maintenance therapy and nonmyeloablative or myeloablative chemotherapy followed by autologous stem cell transplantation [3]. However, the combination of chemotherapy and radiation therapy often results in neurotoxicity. Despite consolidation treatment, relapse is common and the 5-year survival rate remains 30–40% [4–6].

The heterogeneity observed in PCNSL and its microenvironment necessitates the use of individualised treatment strategies. To date, various animal models and cell lines have been used to identify novel therapeutic agents for DLBCL and PCNSL [7,8]. However, only two PCNSL (HKBML, TK) cell lines [9] and a few genuine patient-derived xenografts (PDXs) have been established [10]. These models have certain limitations. For instance, traditional *in vitro* culture requires extended initiation and an exogenous medium to expand tumour cells through passages, which impedes cellular diversity and preservation of key genes [11]. PDX models, which involve direct injection of patient-derived tumour cells into mice, can simulate the characteristics of PCNSL more efficiently; however, skilled surgeons are required for the injection of tumour cells into the mouse brain. Moreover, these models exhibit varying engraftment efficiencies, have limited throughput and require 2–6 months for tumour development [10,12]. With recent advancements, single-cell sequencing has revealed the molecular heterogeneity of tumours and its relationship with unfavourable outcomes [13,14]. In particular, individual tumour cells harbour distinct genetic and epigenetic profiles, which may affect drug resistance and metastasis. Although studies have improved the understanding of recurrent genetic mutations and chromosomal changes [15–18], developing individualised treatment models and strategies for PCNSL remains challenging.

The recent advancements in patient-derived tumour organoids research have effectively addressed the challenges posed by inter-patient and intra-tumoral heterogeneity, establishing a robust and reliable model for scientific investigations [19,20]. These organoids serve as reliable models for drug screening and predictive analysis, facilitating the development of clinical pharmacotherapy. To date, organoids have been used to model various cancers, including glioblastoma [21], neuroendocrine neoplasms [22] and rectal and colorectal cancers [23]. However, the long-term propagation of PCNSL organoids remains unreported. In this study, we designed a robust method for the rapid generation of PCNSL organoids from fresh tumour specimens, eliminating the requirement of single-cell dissociation. Eight patient-derived PCNSL organoids (CLOs) were generated and subjected to comprehensive histological, molecular and genomic analyses. The results validated that the CLOs accurately mimicked inter- and intra-tumoral heterogeneity while retaining many important features of their corresponding parental tumours. In addition, the CLOs demonstrated the potential to predict chemotherapy responses in patients with cancer. These findings highlight the prospective utility of the CLOs in both fundamental and applied research and in evaluating individualised treatment approaches.

## Subjects and Methods

### Human Participants

This study was conducted in accordance with the ethical guidelines established by the Declaration of Helsinki and was approved by the Institutional Review Boards (2022-529) of Huashan Hospital of Fudan University and Fudan University Shanghai Cancer Center (1612167-18). Informed consent was obtained from all participants before tissue collection. Fresh tumour tissues were collected from 9 patients with PCNSL at Huashan Hospital and Fudan University Shanghai Cancer Center between 1^st^ October 2021 and 31^st^ March 2023. All patients were diagnosed with PCNSL according to the 2016 World Health Organization criteria [24].

Table S1 provides a comprehensive overview of the demographic and histological characteristics of each patient. The diagnostic criteria were the same as those described previously [25]. Germinal centre B cell-like (GCB) lymphoma and non-GCB lymphoma were classified using the Hans algorithm [26].

### Treatment and Follow-up

All patients received a methotrexate-based combination immunochemotherapy regimen, which included ibrutinib, dexamethasone and rituximab [27]. The patients were regularly monitored through outpatient visits every 3 months until their demise, ensuring timely updates on their survival status, disease progression and time of death. After treatment, the patients underwent magnetic resonance imaging (MRI) and a clinical follow-up. If clinical signs or symptoms suggested tumour recurrence, MRI was immediately performed to assess the condition of the patients.

### Generation of CLOs from Resected PCNSL Tissues

To form CLOs, we first need to generate host organoids following Lancaster’s method[28]. The process for generating CLO from resected PCNSL tissues is illustrated in Figure 1A. PCNSL tissues were placed in Petri dishes and washed twice with phosphate-buffered saline (PBS; Sigma, #P4474). The tissues were fragmented into small pieces measuring 1–3 mm in diameter, washed once with PBS and introduced into modified DMEM medium supplemented with retinoic acid (RA), and containing ROCK inhibitor (Sigma, #SCM075). Subsequently, the fragmented tissue samples were transferred to 6-well plates and incubated with 1 µL of luciferase-labelled adenovirus and 50 µl of DMEM+RA medium at 37°C for 1 hour. Wild-type organoids (STEMdiff™ Cerebral Organoid Kit) were extracted and positioned on a membrane. A scalpel was used to create an incision measuring approximately 1/3–1/2 of the organoid’s size. After 1 hour of incubation, PCNSL tissues were carefully placed within the incision using forceps. The organoids were fixed using Matrigel (Corning, #354277) and incubated at 37°C for 30 minutes. For culture, the organoids were transferred to 6-well plates, with each well containing 3–4 organoids and 3 mL of DMEM+RA medium, and incubated on a shaker at 37°C overnight. An *in vivo* imaging system (Bio-Real QuikView 2000) was used to detect the signal intensity and size of PCNSL organoids at 2-day intervals over a 14-day culture period.

**Figure 1.**
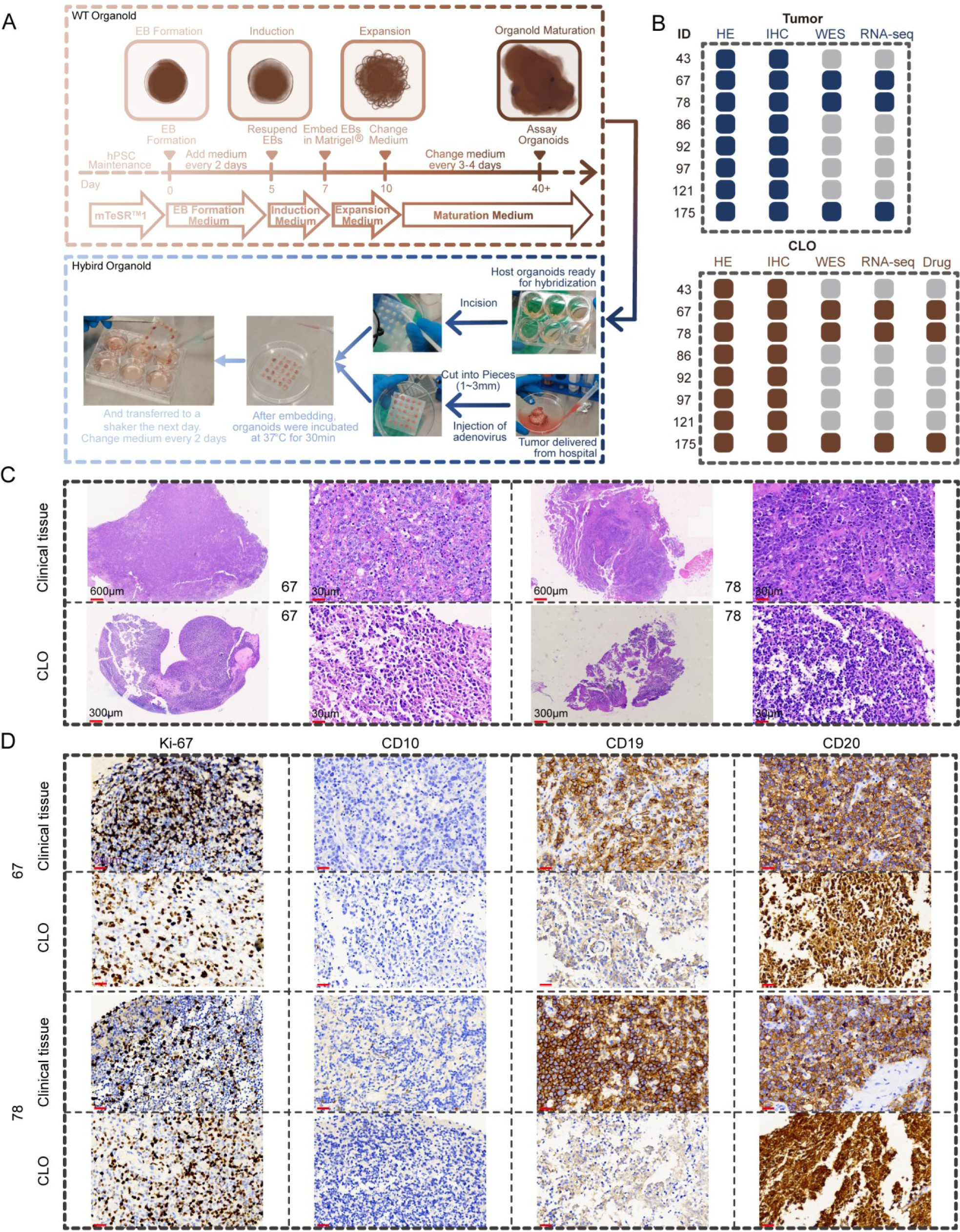
Study Design and Generation of Organoids from Primary Tissue Samples. (A) Diagram Illustrating the Organoid Culture Protocol. (B) A total of 8 organoid lines were successfully established and subjected to comprehensive analyses, including histopathology, whole-exome sequencing, RNA expression profiling, and drug sensitivity assays. DNA from the corresponding primary tumors was also isolated to assess concordance. (C) Comparative H&E Staining of Primary Tumors and Their Derived Organoids. (D) Immunohistochemical Staining for Ki67, CD10, CD19, and CD20 in Parental Clinical Tissues and Corresponding PCNSL Organoids (CLO).

The cooling switch of the instrument was turned on beforehand to decrease the temperature to −80°C. A total of 13.6 µL of D-luciferin (Sigma, #50227) was added to each well 15 minutes before imaging. The 6-well plate was inserted into the instrument 2 minutes in advance, with the camera field of view set at 20 cm × 20 cm and the instrument parameters adjusted to an exposure time of 2 minutes and an EM Gain of 100. After the measurement of signals on day 14, PCNSL organoids were collected for drug sensitivity analysis, RNA sequencing (RNA-seq) and whole-exome sequencing (WES).

### Drug sensitivity analysis

A total of 15 organoids from each patient with PCNSL were individually placed in 24-well plates. Each sample was mixed with 1 mL of DMEM+RA medium supplemented with specific drugs, with drug concentrations being determined based on previous studies (methotrexate [Aladdin, #M408860], 5 μM [29]; ibrutinib [Aladdin, #P127143], 500 nM [30]; dexamethasone [Sigma, #D4902], 300 nM [31]; rituximab [Sigma, #MSQC17], 1 μM [32]). Accordingly, the samples were divided into five groups, with three organoids in each group, as follows: control, methotrexate, ibrutinib, dexamethasone and rituximab. The drug-containing culture medium was replaced every 2 days for the detection of signal intensity. The culture was continued for 20 days, during which organoid samples were collected and frozen for further use. Alterations in the activity of PCNSL organoids treated with specific drugs on different days were examined by analysing their signal intensity and size on the Bio-Real in vivo imaging system. To minimise the impact of human factors on the assessment of signal intensity and size, the average signal intensity and size of the three organoids in each group were calculated. Subsequently, the average signal intensity and size were compared between the experimental and control groups. The growth of PCNSL organoids was evaluated based on the changes in these two parameters.

### Haematoxylin–Eosin and Immunohistochemical Staining

Tumour tissues and CLOs were fixed with 10% paraformaldehyde and embedded in paraffin. The paraffin-embedded tissues were cut into 4-µm-thick sections and stained with haematoxylin–eosin (H&E) using standard protocols. Immunohistochemical (IHC) analysis was performed on the automated Leica BOND-III stainer system using the following antibodies: Ki-67 (ab15580, Abcam), 1:200; CD20 (ab78237, Abcam), 1:200; PAX5 (M701929, DAKO), 1:200; CD19 (ab245235, Abcam), 1:200; CD2 (#9987, Cell Signaling Technology), 1:200; CD10 (ab256494, Abcam), 1:200; BCL2 (#15071, Cell Signaling Technology), 1:200; MUM1 (#62834, Cell Signaling Technology), 1:200; C-MYC (ab32072, Abcam), 1:200; BCL6 (#89369, Cell Signaling Technology), 1:200. IHC staining was evaluated independently by two haematopathologists (J R and ZG X), and discrepancies were resolved by another haematopathologist (JC F).

### TUNEL staining

Tumour tissues and CLOs were fixed with 10% paraformaldehyde and embedded in paraffin. The paraffin-embedded tissues were cut into 4-µm-thick sections. Apoptotic cells were identified via double-immunofluorescence staining with TUNEL (#11684817910, In Situ Cell Death Detection Kit, Roche). The percentage of positive cells was semi-quantitatively calculated by two pathologists independently and was subsequently averaged. Tissue sections were digitally scanned on an KFBIO scanner (KF-PRO-005-EX, Zhejiang, China) and visualised using the KFBIO SlideViewer software.

### Fluorescence in-situ hybridisation

Tumour tissues and CLOs were fixed with 10% paraformaldehyde and embedded in paraffin. The paraffin-embedded tissues were cut into 4-µm-thick sections. Fluorescence in-situ hybridisation (FISH) was performed in accordance with standard protocols using the following commercial probes: MYC break-apart, BCL2 break-apart, IRF4 break-apart and BCL6 break-apart probes (LBP Medicine Science & Technology Co., Ltd, Guangzhou, China). Staining was independently evaluated by two haematopathologists (J R and ZG X), and discrepancies were resolved by another haematopathologist (JC F).

### Whole-exome sequencing

At 2 weeks of CLO culture, at least 2 pieces of organoids from each patient were combined, snap-frozen on dry ice and stored at −80°C until further use. Similarly, parent tumour tissues were frozen and stored until further use. Genomic DNA was extracted from frozen tumour tissues and organoids using the QIAamp DNA FFPE Tissue Kit according to the manufacturer’s protocol (QIAGEN). The quality of extracted DNA was estimated using the Qubit high-sensitivity dsDNA assay kit (Thermo Fisher Scientific) and Nanodrop 2000 spectrophotometer (Thermo). A total of 250 ng of extracted DNA was fragmented to the target average size of 150–200 bp using the Covaris LE220 Sonicator (Covaris), and DNA libraries were prepared using the KAPA Hyper Prep kit (Roche). The 3′ and 5′ overhangs of the fragmented DNA were repaired using End-Repair Mix (a component of SureSelect XT HS2) and purified using Agencourt AMPure XP Beads (Beckman). ‘A’ tails were added to the purified fragments using A-tailing Mix (a component of SureSelect XT HS2), and the fragments were ligated with adapters using a DNA ligase (a component of Sureselect XT HS2). The adapter-ligated DNA fragments were amplified using Herculase II Fusion DNA Polymerase (Agilent), and pre-captured libraries containing exome sequences were detected using SureSelect HS Human All Exon V8 (Agilent). The prepared libraries were quantified using the Qubit 3.0 Fluorometer dsDNA HS Assay Kit (Thermo Fisher Scientific), and their size distribution was analysed on the Agilent Bioanalyzer 4200 system (Agilent).

The quality of DNA-sequencing data was controlled using FastQC (v0.11.8). Adapters were trimmed using fastp (v0.12.5). QC statistics were recorded and summarised using Qualimap (v2.2.1). Data cleanup and variant detection were performed using Genome Analysis Toolkit (GATK, version v4.1.9.0.) according to the standard practice guidelines. The reads were aligned to the reference genome of GRCh37 (kindly provided by the Broad Institute) using BWAMEM [33] with default parameters on the GATK public FTP server. Pre-processing of data, including marking of PCR duplicates and base quality recalibration, was performed using Picard (v.1.141) and GATK (v4.1.9.0) according to standard protocols. The aligned sequencing data were locally realigned to improve sensitivity and reduce artifacts before the identification of single-nucleotide variants (SNVs). In addition, insertions and deletions (INDELs) were detected using GATK (v4.1.9.0) [34]. Somatic SNVs and INDELs were identified using MuTect2 (v1.1.4), and consensus calls were used for the final analysis. GATK FilterMutectCalls was used to remove unreliable somatic mutations. Somatic copy number variants (CNVs) were identified based on segmented copy number profiles generated from WES using xHHM (v6.1). Point mutations were decomposed into several different mutational features using non-negative matrix factorisation [35]; subsequently, these features were clustered with 78 known mutation features from the COSMIC website (http://cancer.sanger.ac.uk/cosmic/signatures).

### RNA-seq

At 2 weeks, at least 5 pieces of organoids from each patient were combined, snap-frozen on dry ice and stored at −80℃ until further use. Similarly, parent tumour tissues were frozen and stored until further use. Total RNA was isolated from frozen tumour tissues and organoids using the RNeasy mini kit (Qiagen, Germany). The concentration and quality of extracted RNA were determined using Qubit®3.0 Fluorometer (Life Technologies, USA) and Nanodrop One spectrophotometer (Thermo Fisher Scientific Inc., USA). The integrity of extracted RNA was assessed using Agilent 2100 Bioanalyzer (Agilent Technologies Inc., USA), and samples with an RNA integrity number (RIN) of >7.0 were used for sequencing. A total of 1 µg of RNA was used as input material, and paired-end libraries were synthesised using the mRNA-seq Lib Prep Kit for Illumina (ABclonal, China) following the sample preparation guide. Purified libraries were quantified using Qubit® 2.0 Fluorometer (Life Technologies, USA) and validated using Agilent 2100 Bioanalyzer (Agilent Technologies, USA) to confirm the insert size and calculate the molar concentration. The libraries were diluted to 10 pM, clustered was using cBot and sequenced on the Illumina NovaSeq 6000 platform (Illumina, USA).

The imaging data captured by high-throughput sequencers were converted to sequencing data (reads) using CASAVA (base calling). Quality assessment and pre-processing of the sequencing data were performed using the FastQC software (version 0.12.1). To ensure the quality and reliability of data analysis, the fastp software [36] was used to filter the sequencing data as follows: removing adapter-containing reads, trimming bases with a quality score (Q) of <20 at the 3′-end, removing reads shorter than 25 bases and eliminating ribosomal RNA reads from target species. Additionally, Q20, Q30 and GC content were calculated to obtain clean data. All subsequent analyses were performed using high-quality clean data.

The reference genome (GRCh38.108) and gene model annotation files were downloaded from the genome website. HISAT2 (v2.1.0) was used to build the reference genome index and align paired-end clean reads to the reference genome [37]. FeatureCounts (1.5.0-p3) was used to count reads mapped to each gene to quantify gene expression [38]. The gene expression matrix of all samples was compiled [39], and fragments per kilobase of transcript per million mapped reads (FPKM) values were calculated for each gene based on its length [40]. The DESeq2 software (1.20.0) was used for differential expression analysis between two groups [41]. The obtained P-values (padj) were adjusted using the Benjamini–Hochberg method to control the false discovery rate. A padj value of ≤0.05 and |log2(foldchange)| value of ≥1 indicated significant differential gene expression. Principal component analysis (PCA) was performed based on FPKM values using the ggplot2 package in R (version 3.5.0). The distribution of differentially expressed genes across groups was visualised on volcano plots generated using the ggplot2 package in R (version 3.5.0). Pearson correlation analysis was performed using the cor() function in R (version 3.5.0, ggplot2 package). Hierarchical clustering was visualised on heatmaps generated using the pheatmap package in R (version 3.5.0, gglpot2 package). KEGG pathway enrichment analysis of differentially expressed genes was performed using the clusterProfiler (3.8.1) software [42].

### Statistical analysis

Statistical analysis was performed using the GraphPad Prism 9 software (GraphPad Software, La Jolla, CA, USA). Comprehensive details regarding the statistical tests and sample sizes are provided in Figure Legends. All data were expressed as the mean ± SE/SD. For the quantification of markers in IHC and immunofluorescence analyses, the samples being compared were processed simultaneously and captured using consistent imaging settings. Positively stained cells were manually counted using the Cell Counter function in the ImageJ software (NIH). For each set of quantification, images were selected from the peripheral regions of three organoids. All cell counts were statistically evaluated using one-way ANOVA. Statistical significance was indicated as follows: ***, *p* < 0.001; **, *p* < 0.01; *, *p* < 0.05.

## Results

### Successful Establishment and Histological Confirmation of PCNSL-derived Organoids

Surgically resected or biopsy-obtained tissue samples were collected from 9 patients newly diagnosed with PCNSL, with 6 samples from Huashan Hospital and 3 from Shanghai Cancer Center. The baseline characteristics of all patients are provided in Table S1. The tissues were collected from treatment-naive patients at the time of CLO generation. We developed a non-invasive method to culture CLOs using small tumour tissue fragments, preserving the tissue architecture and cell–cell interactions of the original tumours and minimising clonal selection. Organoids were typically formed within 2 weeks (Figure 1A). The number of organoids generated varied based on the tissue size; in particular, surgical samples produced 20–30 primary organoids, whereas biopsy samples produced 3–7 organoids. Of the 9 samples, 8 samples successfully yielded continuously proliferating organoid cultures. Although the remaining sample could produce organoids, they failed to grow in vitro. The organoids generated from the 8 samples were subjected to comprehensive histological, molecular, genomic and drug sensitivity analyses (Figure 1B). Bright-field images were captured to assess the temporal progression of organoid development (Figure S1A).

Histological analysis was performed to examine whether CLOs mimicked the characteristics of parental tumour tissues. Each CLO was rigorously examined by an experienced neuropathologist, who confirmed the retention of hallmark PCNSL features. Consistently, H&E staining showed that the CLOs exhibited heterogeneous morphological features similar to those observed in parental tumour tissues (Figure 1C, Figure S1B). Both CLOs and parental tumour tissues exhibited diffuse cell proliferation characterised by the presence of medium-to-large cells with a round or oval shape, prominent nucleoli, scant cytoplasm, basophilic hues and pathological mitotic features. Notably, the CLOs had a slightly reduced cell size, subtle chromatin pyknosis and minor degeneration in tumour cells. In addition, Ki-67 immunostaining showed that the CLOs preserved the proliferative index of parental tumour tissue (Figure 1D, Table S2, Table S3).

To identify the cell subsets within the organoids, immunohistological analysis was performed using a targeted panel of markers, including CD10, CD19 and CD20. Quantitative analysis revealed a high degree of concordance in the expression of CD10, CD19 and CD20 between parental tumour tissues and their respective CLOs (Figure 1D, Table S2, Table S3). The histological features and expression characteristics of the parental tumor markers are fully preserved by CLOs.

### Genomic Analysis Validated the Homogeneity Between CLOs and Parental Tumours

To determine whether CLOs maintain genomic alterations found in parental tumors, WES was performed to assess the genomic characteristics of the eight matched pairs of parental tumours and their corresponding CLOs. We focused on somatic variants listed in a recent comprehensive study of PCNSL genomics[15]. Genome-wide analysis of CNVs revealed that the CLOs largely retained the copy number deletions and amplifications observed in parental tumour tissues (Figure 2A). A high degree of concordance was observed in PCNSL in the CNVs of cancer-associated genes between the CLOs and parental tumour tissues, with an average similarity of 80.11% and closely aligned copy number ratios (Figure 2B, Figure S2A). On comparing SNVs in PCNSL-related genes between parental tumour tissues and their corresponding CLOs, we found that the CLOs contained an average of 81.85% of the selected cancer-associated genes from parental tumour tissues (Figure 2C, Figure S2B). Notably, mutations in key PCNSL-associated genes, such as PIM1, MYD88 and CD79B, were highly preserved across most CLO lines. Additionally, the CLOs maintained the overall mutational burden and the spectrum of point mutations observed in parental tumour tissues (Figures 2D, Figure S2C). The mutational signatures exhibited a similarity of 91% between the CLOs and parental tumour tissues, with the similarity between individual CLOs and their corresponding parental tumours ranging from 74% to 98% (Figure 2D). Altogether, these results indicate the high genomic fidelity of CLOs, establishing them as a robust model for further research on PCNSL.

**Figure 2.**
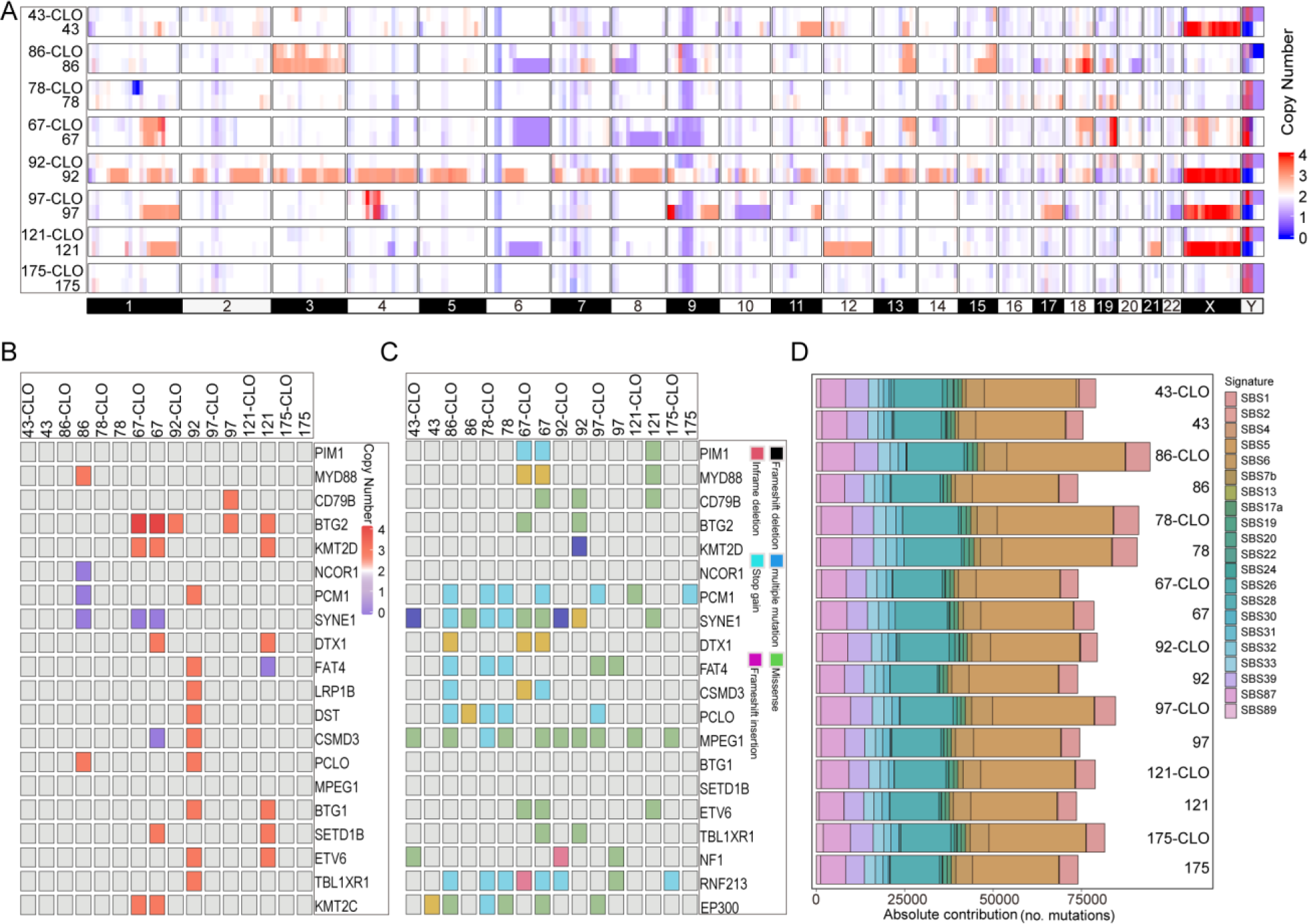
Genetic Fidelity of PCNSL organoid (CLO) Mirrors Corresponding Parental Tumors. (A) Comparative Analysis of Somatic Copy Number Alterations Across 8 Paired CLOs and Parental Tumors. (B) Heatmap Illustrating the Concordance in Altered Cancer Gene Copy Numbers Between CLOs and Parental Tumors. (C) Heatmap Depicting the Preservation of Somatic Mutations in Cancer-Related Genes Across CLOs and Parental Tumors. (D) Stacked Bar Chart Showcasing the Proportional Distribution of COSMIC Mutational Signatures in Both CLOs and Parental Tumors.

### Transcriptomic Analysis Highlights Similar Gene Expression Profiles Between CLOs and Parental Tumours

To compare the gene expression profiles of parental tumour tissues and their corresponding CLOs, transcriptomic analysis was performed using the bulk RNA-seq data of three matched pairs of parental tumour tissues and CLOs that were cultured for 2 weeks. As shown in the heatmap in Figure 3A, a high degree of similarity was observed between the gene expression profiles of CLOs and parental tumour tissues, with transcriptome-wide comparisons revealing a robust correlation (r > 0.75). PCA revealed the intrinsic heterogeneity between the CLOs and parental tumour tissues (Figure 3B). Despite the pronounced inter-tumoral heterogeneity of PCNSL, the gene expression patterns of individual CLOs were consistent with those of their respective parental tumour tissues (Figure 3C). KEGG pathway analysis validated the conservation of the parental gene expression patterns in CLOs (Figure 3D).

**Figure 3.**
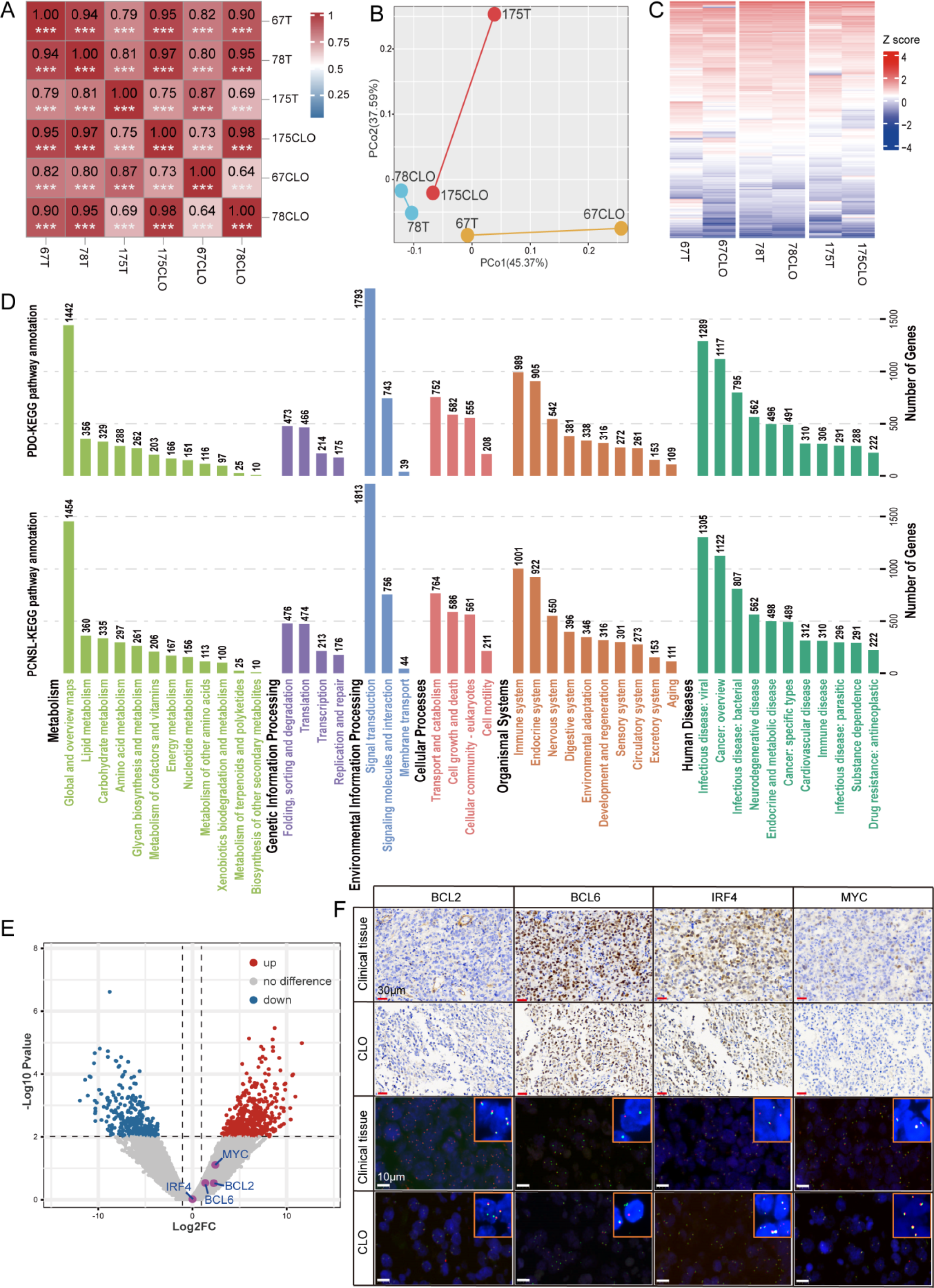
Preservation of Inter- and Intra-Tumoral Gene Expression Heterogeneity in PCNSL organoid (CLO) Relative to Parental Tumors. (A) Heatmap illustrating transcriptome-wide Pearson correlations between gene expression in parental tumors and their corresponding CLOs, as assessed by RNA-seq. (B) Principal Component Analysis (PCA) plots of RNA-seq gene expression data, comparing subregional samples from parental tumors and their respective CLOs. (C) Heatmap showcasing the top 10,000 most variably expressed genes in parental tumors. (D) KEGG pathway analysis to evaluate the conservation of gene expression pathways between parental tumors and CLOs. (E) Volcano plot highlighting differentially expressed genes in CLOs relative to their corresponding parental tumors. (F) Representative images of immunostaining and Fluorescence In Situ Hybridization (FISH) in both the parental tumors and their corresponding CLOs. ***: *P*<0.001.

As shown in Figure 3E, volcano plots did not reveal significant changes in the expression of key PCNSL markers, including BCL6, BCL2, MYC and IRF4, between the CLOs and parental tumour tissues. This finding was corroborated by immunohistological analysis (Figure 3F, Tables S2–S3). The rearrangement statuses of BCL6, BCL2, MYC and IRF4 were consistent between the CLOs and parental tumour tissues (Figure 3F, Table S4). Altogether, these results validated that the CLOs closely replicated the genetic landscape of their respective parental tumours, demonstrating their utility as clinically relevant models.

### Response of CLOs to Chemotherapeutic and Targeted Drugs

In clinical practice, the efficacy of individual drugs in a certain combination therapy is often obscured in most patients. Organoid models can be used to evaluate the effectiveness of individual drugs. In this study, therapeutic profiling was performed on three CLOs using four drugs. The clinical information of the three patients (IDs: 67, 78, 175) is provided in Table S1. To validate the *in vitro* responsiveness of CLOs to ibrutinib, the signal intensity (Figure 4A) and size (Figure S3A) of all three CLOs were examined. The signal intensity curves indicated that the CLOs from patients 67, 78 and 175 were sensitive to ibrutinib (Figure 4A). This finding was corroborated by the size–response curves of the three CLOs (Figure S3A). In addition, the three CLOs were sensitive to methotrexate (Figures 4B and S3B) but unsensitive to dexamethasone (Figures 4C and S3C) and rituximab (Figures 4D and S3D). Tables S5 and S6 contain the raw data pertaining to the images and curves that depict variations in signal intensity and size of individual organoids.

**Figure 4.**
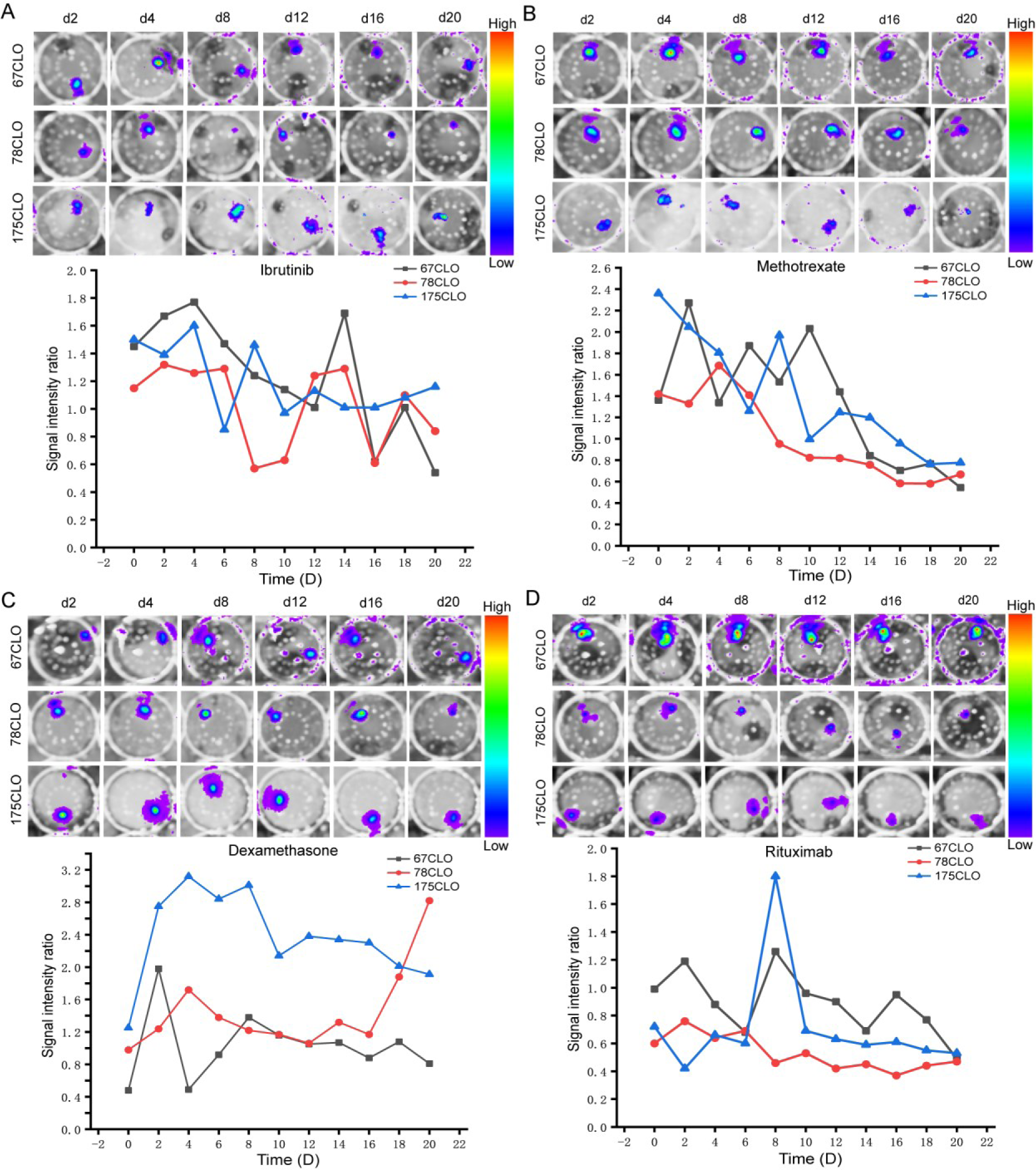
Evaluating PCNSL organoid (CLO) Responses to Ibrutinib, Methotrexate, Dexamethasone, and Rituximab Treatments. (A) Time-course Analysis of Organoid Sensitivity to Ibrutinib in Three PCNSL Patients from Day 2 to Day 20; Mean Signal Intensity Ratios ± SE from Three Independent Experiments. (B) Time-course Analysis of Organoid Sensitivity to Methotrexate in Three PCNSL Patients from Day 2 to Day 20; Mean Signal Intensity Ratios ± SE from Three Independent Experiments. (C) Time-course Analysis of Organoid Sensitivity to Dexamethasone in Three PCNSL Patients from Day 2 to Day 20; Mean Signal Intensity Ratios ± SE from Three Independent Experiments. (D) Time-course Analysis of Organoid Sensitivity to Rituximab in Three PCNSL Patients from Day 2 to Day 20; Mean Signal Intensity Ratios ± SE from Three Independent Experiments.

Furthermore, the responsiveness of CLOs to key therapeutic agents—methotrexate, ibrutinib, dexamethasone and rituximab—was evaluated using in situ cell death detection assays and Ki-67 immunostaining. As shown in Figures 5A, 5B and Figures S4, the proportion of TUNEL-positive cells in the three CLOs was significantly higher in the methotrexate and ibrutinib treatment groups than that in the control group. On the contrary, the proportion of TUNEL-positive cells in the three CLOs was similar between the dexamethasone or rituximab treatment group and the control group (Figures 5A, 5B, and Figures S4). Ki67 immunostaining revealed that the proportion of Ki67-positive cells in the three CLOs was higher in the methotrexate and ibrutinib treatment groups than that in the control group, whereas it was comparable between the dexamethasone or rituximab treatment group and the control group (Figure 5C, 5D). The statement indicates that CLOs can effectively capture the variations in clinical drug responses observed among different parent tumors.

**Figure 5.**
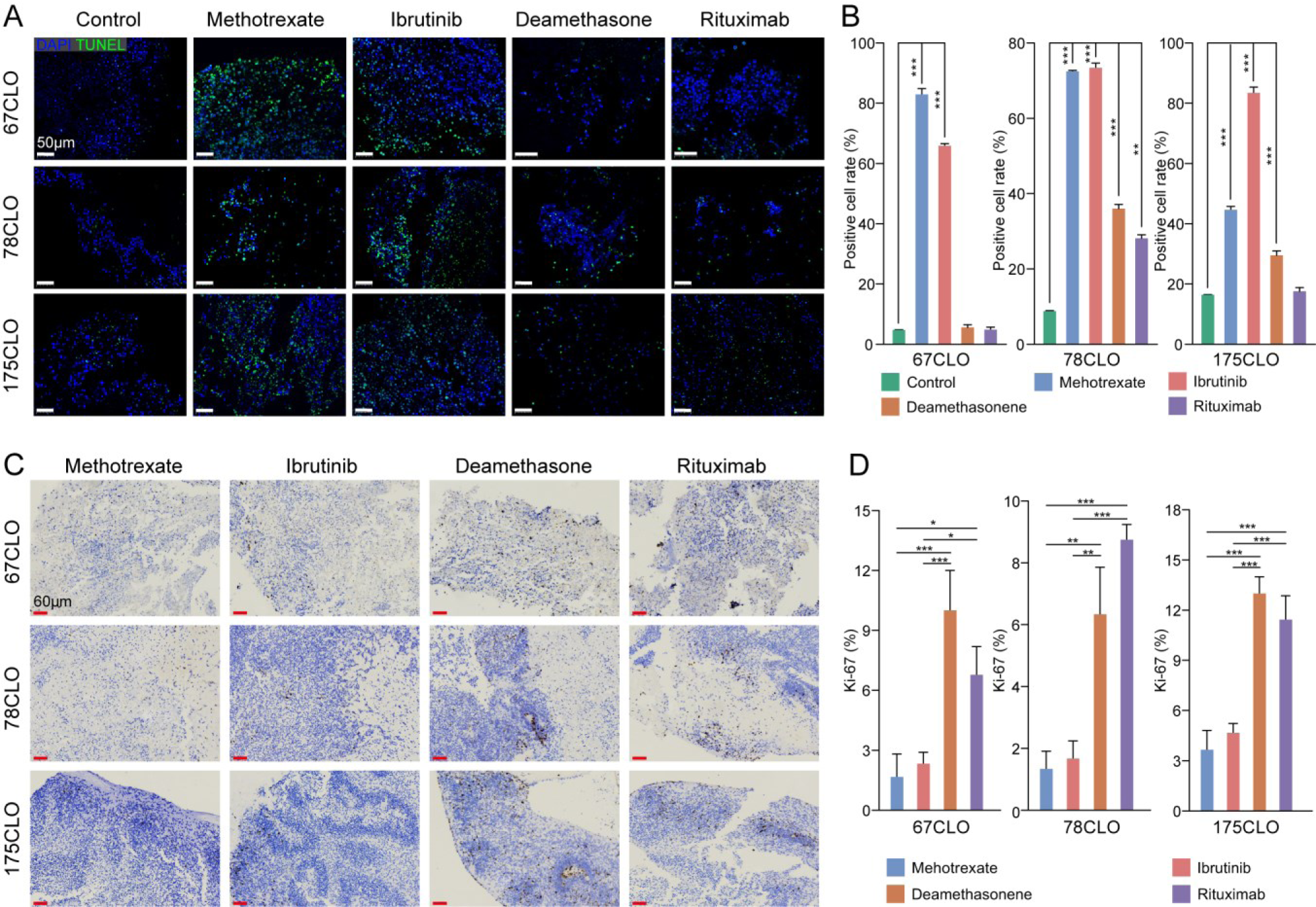
Assessing Drug Sensitivity in PCNSL organoid (CLO) via TUNEL and Ki-67 Assays. (A) TUNEL Assay Outcomes for Ibrutinib, Methotrexate, Dexamethasone, and Rituximab Treatment Groups. (B) Quantitative Analysis of TUNEL-Positive Cells Across Various Treatments. (C) Ki-67 Immunostaining Results for Ibrutinib, Methotrexate, Dexamethasone, and Rituximab Treatment Groups. (D) Quantitative Enumeration of Ki-67-Positive Cells Across Treatment Conditions. For each data set, images were sourced from the peripheral regions of three distinct organoids. All cell count metrics were statistically analyzed using one-way ANOVA. Statistical significance levels are denoted as follows: \*\*\**P* < 0.001, \*\**P* < 0.01, \**P* < 0.05, and deemed not statistically significant when *P* > 0.05.

### Alignment of Clinical Outcomes with Drug Sensitivity in Tumour Organoids

In addition to in vitro assays, MRI validated the correlation between the drug sensitivity of CLOs and actual patient outcomes (Figure 6A). In particular, patients whose corresponding CLOs exhibited significant sensitivity to both ibrutinib and methotrexate showed marked improvements in responsiveness to these drugs in clinical settings. These improvements were quantified based on radiological enhancements observed on MRI, demonstrating the utility of the CLOs in determining effective chemotherapy regimens (Figure 6B). Altogether, these results suggest that the CLOs can effectively predict the actual drug responses of patients in clinical settings.

**Figure 6.**
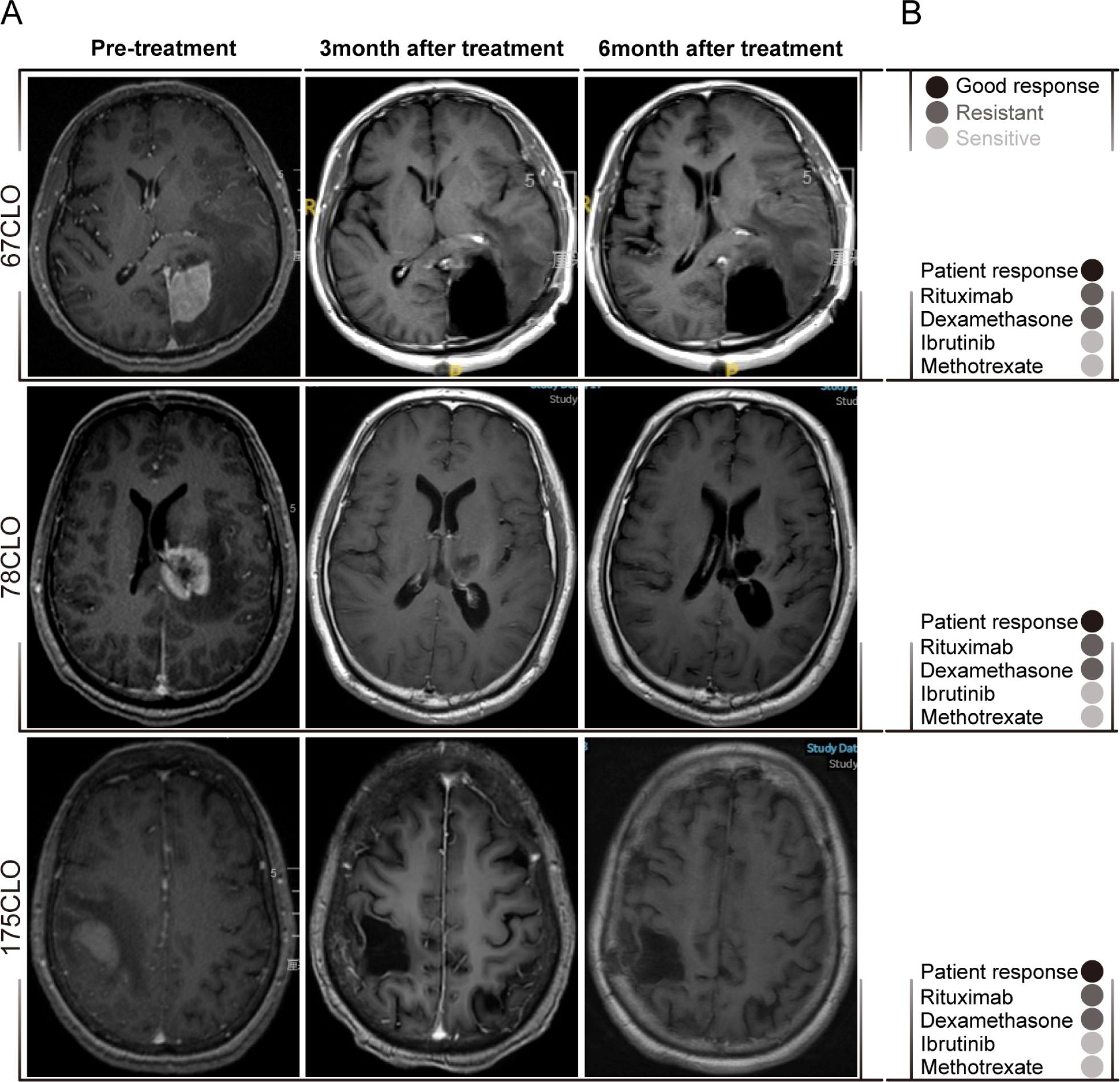
Clinical Outcomes in Patients Align with Drug Sensitivity in Tumor Organoids. (A) Clinical Outcome Assessment: MRI Images of Patient 67, 78, 175 Treated with Ibrutinib, Methotrexate, Dexamethasone, and Rituximab, Captured at 3 months after treatment, 6 months after treatment, and Pre-treatment. (B) Correlation Between Organoid Response Data and Clinical Outcomes in Patients.

## Discussion

In contrast to previously reported PCNSL cell lines and PDX models, the CLOs developed in this study preserve native cell–cell interactions. The limited availability of patients with PCNSL poses challenges to the development of pre-clinical models. To overcome this challenge, we collected PCNSL specimens to establish organoids. The CLOs developed in this study accurately simulated the heterogeneity of their corresponding parental tumours as evidenced by the following findings: (1) Histological analysis revealed similar tissue architecture and cellular morphologies; (2) IHC staining indicated the presence and continual generation of a similar spectrum of cell types; (3) RNA sequencing indicated the maintenance of similar transcriptomic signatures; (4) WES validated the preservation of SNVs and CNVs at similar frequencies; (5) Drug sensitivity analysis revealed a high degree of concordance between chemotherapy responses in patients and CLOs.

The method used to generate CLOs with diverse mutational profiles in this study is both robust and rapid and serves as a valuable resource for future biological studies, therapeutic testing and clinical decision-making. An essential aspect of this method is the preservation of native cell–cell interactions, which facilitates the formation of patient-derived organoids through a fully defined culture system[43], without the requirement of single-cell dissociation. This approach may contribute to the maintenance of biological properties similar to those of parental tumours. In this study, the CLOs and their corresponding parental tumours had a comparable proportion of actively proliferating cells and similar heterogeneous morphologies. Organoid biobanks have been established for various types of cancers such as glioblastoma, pancreatic cancer, liver cancer, neuroendocrine neoplasms, prostate cancer, breast cancer, bladder cancer, ovarian cancer and gastrointestinal cancer. However, such biobanks have not yet been established for PCNSL. In this study, we developed eight CLOs for PCNSL and characterised them via histological analysis, RNA-seq, WES and drug sensitivity analysis. Given that the collection of tumour tissues and generation of CLOs is an ongoing process, the biobank developed in this study serves as a valuable resource for future biological investigations and therapeutic testing related to PCNSL.

Developing algorithms that can accurately predict drug sensitivity based on the unique genetic profiles of patients with cancer remains challenging. At present, organoids offer several advantages in drug development, cancer screening and prediction of therapeutic responses [44]. Patient-derived organoids have been demonstrated to effectively predict drug responses in vitro [45] [46]. For example, Yao *et al.* [47] reported that the responsiveness of patients with rectal cancer to chemoradiation was highly similar to that of the corresponding organoids, with an accuracy of 84.43%, sensitivity of 78.01% and specificity of 91.97%. In this study, we evaluated the responsiveness of CLOs to four therapeutic agents for PCNSL, namely, methotrexate, ibrutinib, dexamethasone and rituximab. The results showed that the CLOs were markedly sensitive to methotrexate and ibrutinib and resistant to dexamethasone and rituximab. Subsequently, patients with PCNSL, from whom tumour tissues were collected to develop the CLOs, were treated with the aforementioned pharmacological agents. Therapeutic efficacy was evaluated through quantitative MRI-based assessment of tumour volumetrics. The clinical outcomes exhibited a significant concordance with the in vitro drug sensitivity profiles, resulting in favourable prognostic outcomes without disease recurrence during the defined observational period. The consistency between the drug responsiveness of CLOs and the corresponding clinical efficacy of drugs in patients with PCNSL strongly highlights the potential of CLOs as an important tool in the development of individualised therapeutic strategies for PCNSL.

In this study, CLOs exhibited heightened sensitivity to methotrexate and ibrutinib, while displaying a comparatively diminished response to dexamethasone and rituximab. Dexamethasone primarily addresses hematological malignancies by inducing lymphocyte apoptosis[48]. Typically, it is administered in conjunction with chemotherapy agents to augment tumor cell eradication; however, its efficacy may be constrained when employed as a standalone treatment. Consequently, the utilization of dexamethasone alone appears to lack effectiveness for CLOs. Rituximab, a targeted therapy, functions by specifically attaching to the CD20 molecule present on the outer membrane of B lymphocytes, thereby inducing cytotoxic effects via antibody-dependent and complement-dependent mechanisms[49]. Our findings align with previous clinical investigations, which have failed to demonstrate substantial therapeutic benefits of rituximab in the treatment of PCNSL[50].

Although the CLOs simulate many features of their corresponding parental tumours, they have certain limitations: (1) The primary treatment method for PCNSL is MTX-based combination immunochemotherapy, with surgery not being a preferred option. The collection of optimal tumour tissues depends on CT- or MRI-guided stereotactic biopsy, which may result in a limited amount of biopsy tissue. Furthermore, a significant portion of the biopsy tissue is required for neuropathology to confirm the diagnosis. A low amount of tissue frequently results in an inadequate quantity of organoids for fundamental characterisation and targeted therapy testing. Therefore, the amount of accessible tumour tissue plays an important role in the production of CLOs. In this study, of the 8 CLOs, only 3 CLOs were viable for targeted therapy testing. (2) The comprehensive characterisation of parental tumours and CLOs, as outlined in this study, may present challenges owing to differences in the initial tumour volume and the rate of expansion of CLOs. A global collaborative network may expand the collection of PCNSL organoids, thereby surpassing the existing constraints owing to the infrequency of PCNSL. (3) Owing to the presence of the blood-brain barrier, sensitive drugs undergoing organ filtration might not be capable of reaching intracranial sites to exert their pharmacological effects.

In conclusion, the patient-derived organoids developed in this study effectively mimic the heterogeneity and essential characteristics of PCNSL and hence serve as a promising tool for prompt evaluation of individualised treatment responses and extensive investigation in fundamental and applied research on PCNSL.

## Availability of data and materials

The WES and RNA-seq raw sequence data have been deposited in the Genome Sequence Archive[51] in National Genomics Data Center[52], China National Center for Bioinformation / Beijing Institute of Genomics, Chinese Academy of Sciences (GSA-Human: HRA005755; https://ngdc.cncb.ac.cn/gsa-human).

Any additional information required to reanalyze the data reported is available from the lead contact upon request.

## Authors’ contributions

WJ C, and SJ L conceptualized and designed this study. SJ L, YZ L, JN W, and JR performed most experiments. ZG X, CX L, and SJ L performed partial experiments. SJ L, CX L, and JR finished the acquisition and analysis of data. SJ L, YZ L, and JN W prepared figures, performed the statistical analysis, and wrote original draft. WJ C, and SJ L reviewed and supervised the manuscript. All authors read and approved the final manuscript

## Acknowledgement

The authors extend their gratitude to Yiyin Zhang from KingMed Diagnostics in Shanghai for his invaluable assistance with data processing. They also wish to express their sincere thanks to the colleagues at Huashan Hospital of Fudan University and Fudan University Shanghai Cancer Center for their crucial support during the specimen collection process. Appreciation is also extended to ShanghaiTech University for providing the research platform.

