## Supplemental tables and figures for "Establishment and Molecular Characterization of Patient-Derived Organoids for Primary Central Nervous System Lymphoma"

Table S1. PCNSL patients' baseline characteristic

| No. | Sex | Age | COO Class | HIV | EB | Derived of tissue | Treatment |
|---|---|---|---|---|---|---|---|
| 43 | male | 58 | non-GCB | - | - | aspiration biopsy | HD-MTX based chemotherapy |
| 67 | male | 67 | non-GCB | - | - | surgery | HD-MTX based chemotherapy |
| 78 | male | 51 | non-GCB | - | - | surgery | HD-MTX based chemotherapy |
| 86 | female | 75 | GCB | - | - | aspiration biopsy | HD-MTX based chemotherapy |
| 92 | female | 55 | non-GCB | - | - | aspiration biopsy | HD-MTX based chemotherapy |
| 97 | male | 75 | non-GCB | - | - | aspiration biopsy | HD-MTX based chemotherapy |
| 121 | female | 58 | non-GCB | - | - | aspiration biopsy | HD-MTX based chemotherapy |
| 175 | male | 53 | non-GCB | - | - | surgery | HD-MTX based chemotherapy |

Table S2. PCNSL patients' histopathological characterization

| No. | ki67 | CD20 | CD19 | CD10 | BCL2 | MUM1 | BCL6 | C-MYC |
|---|---|---|---|---|---|---|---|---|
| 43 | 85% | + | + | - | + | + | + | 50% |
| 67 | 70% | + | + | - | + | + | + | 70% |
| 78 | 55% | + | + | - | + | + | + | 50% |
| 86 | 35% | + | + | - | + | - | + | 60% |
| 92 | 90% | + | + | - | - | + | + | 30% |
| 97 | 90% | + | + | - | + | + | + | 70% |
| 121 | 80% | + | + | - | + | + | + | 40% |
| 175 | 80% | + | + | - | + | + | + | 60% |

Table S3. CLO' histopathological characterization

| No. | Ki-67 | CD20 | CD19 | CD10 | BCL2 | MUM1 | BCL6 | C-MYC |
|---|---|---|---|---|---|---|---|---|
| 43 | 30% | + | + | - | + | + | + | 30% |
| 67 | 20% | + | + | - | + | + | + | 70% |
| 78 | 20% | + | + | - | + | + | + | 40% |
| 86 | 15% | + | + | - | + | - | + | 60% |
| 92 | 50% | + | + | - | - | + | + | 40% |
| 97 | 40% | + | + | - | + | + | + | 50% |
| 121 | 35% | + | + | - | + | + | + | 30% |
| 175 | 45% | + | + | - | + | + | + | 40% |

Table S4. PCNSL patients' and CLO' gene rearrangement statuses

| | PCNSL | | | | | CLO | | | |
|---|---|---|---|---|---|---|---|---|---|
| No. | BCL2 | IRF4 | BCL6 | MYC | | BCL2 | MUM1 | BCL6 | MYC |
| 43 | + | + | + | + | | + | + | + | - |
| 67 | + | + | + | + | | + | + | + | + |
| 78 | + | + | - | - | | + | + | - | - |
| 86 | + | - | + | + | | + | - | + | + |
| 92 | - | + | + | - | | - | + | + | - |
| 97 | + | + | + | + | | + | + | + | + |
| 121 | + | - | + | - | | + | - | + | - |
| 175 | - | + | + | + | | - | + | + | + |

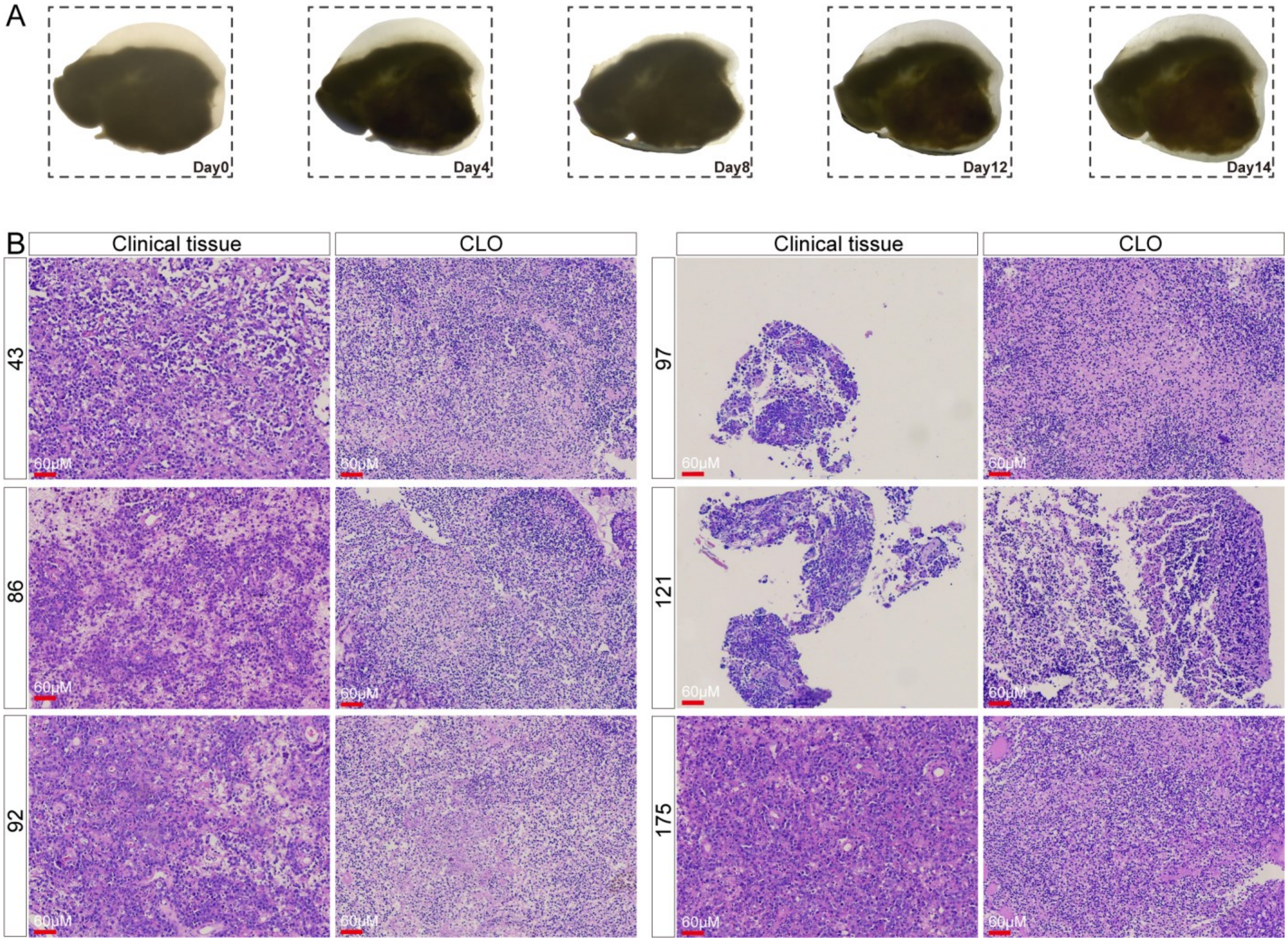


**Figure S1.** Histomorphology of PCNSL organoid.

(A) Temporal Progression of Organoid Development Captured in Sample Bright-Field Images.

(B) Histopathological Characterization of Organoids from Primary Tissue Samples. Comparative H&E Staining of Primary Tumors and Their Derived Organoids.

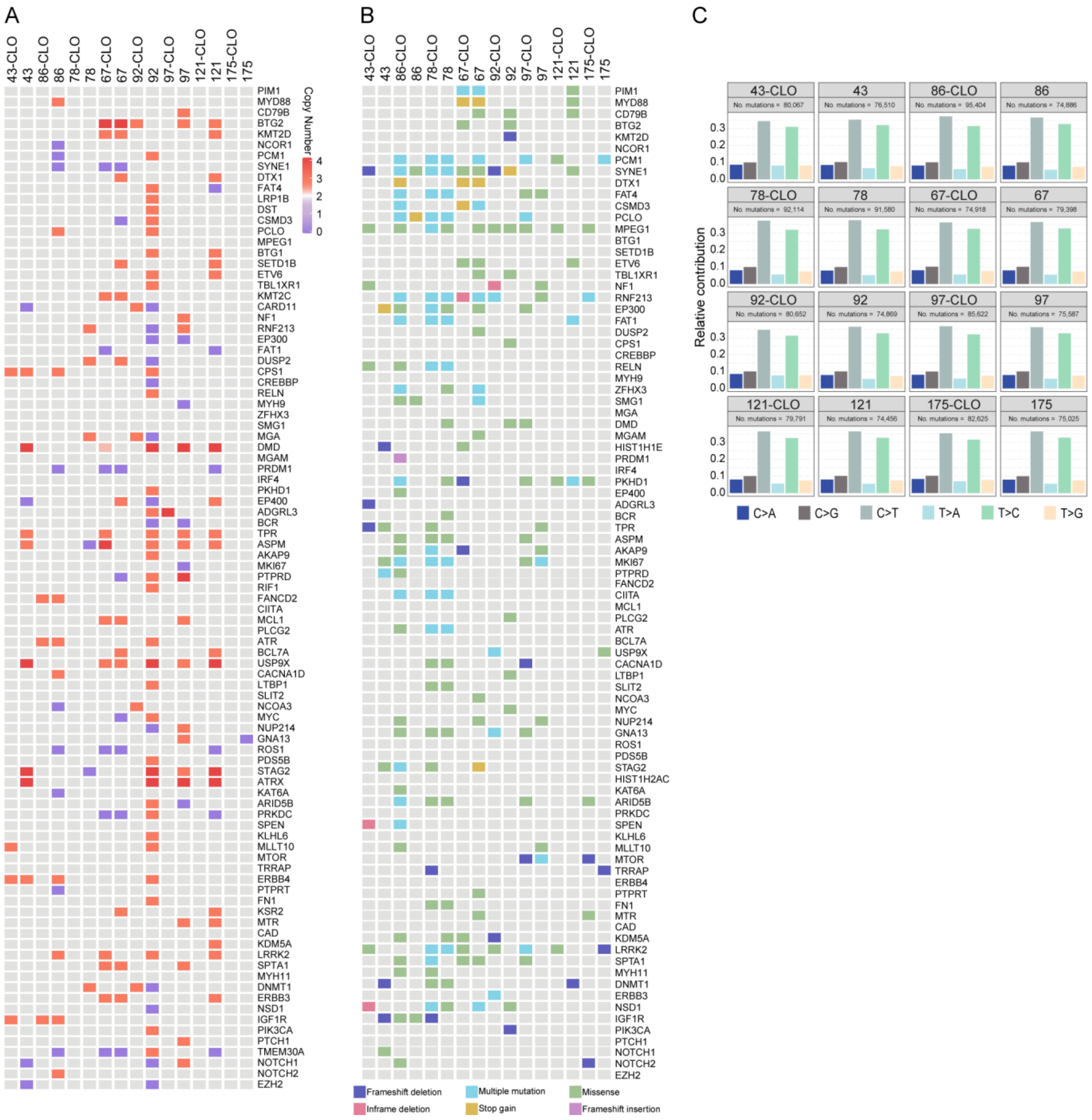


**Figure S2.** Genetic Fidelity of PCNSL organoid (CLO) Mirrors Corresponding Parental Tumors.

(A) Heatmap Illustrating the Concordance in Altered Cancer Gene Copy Numbers Between CLOs and Parental Tumors.

(B) Heatmap Depicting the Preservation of Somatic Mutations in Cancer-Related Genes Across CLOs and Parental Tumors.

(C) Bar Graphs Highlighting the Consistency in Point Mutation Frequencies Between Matched Tumor Tissues and CLOs.

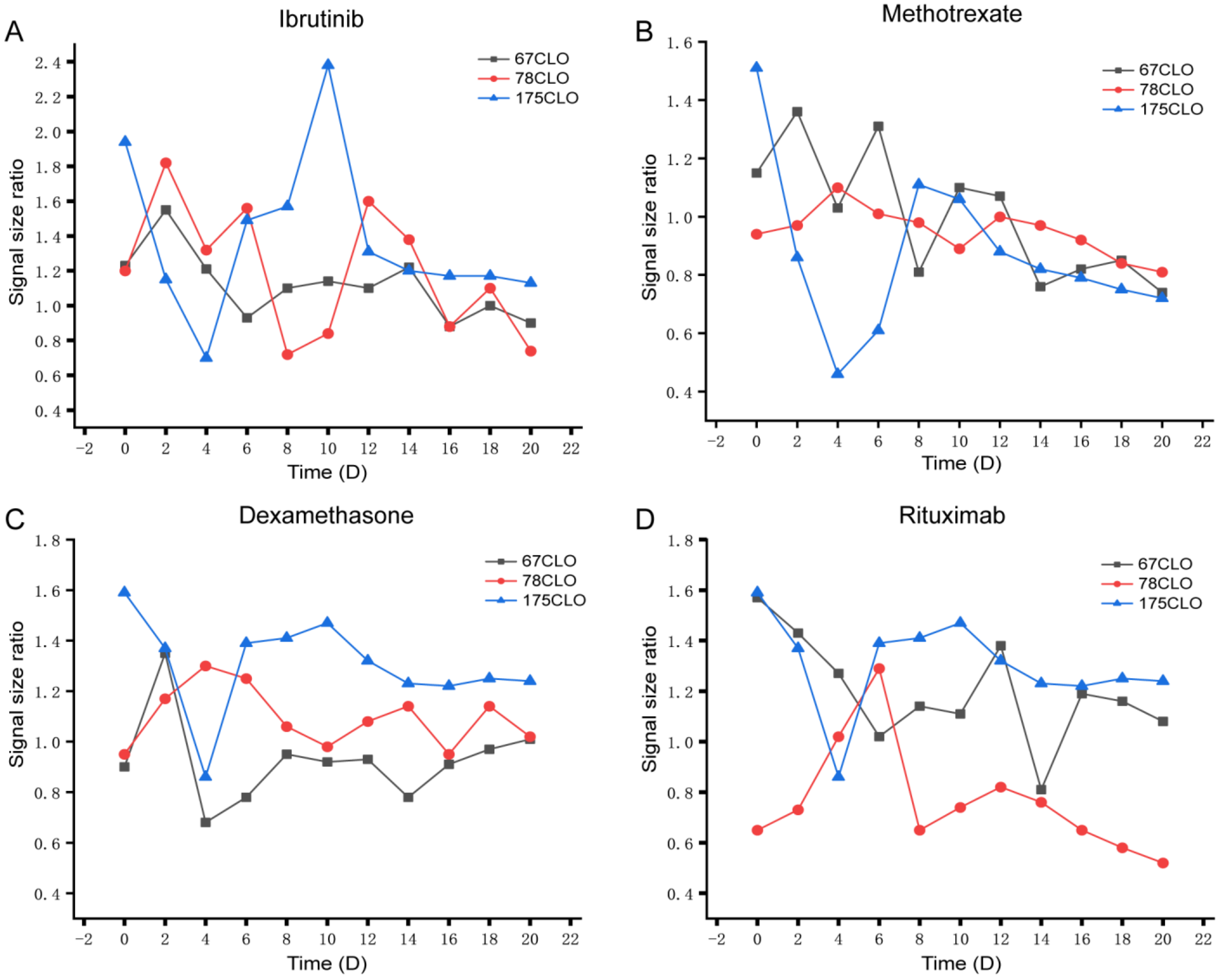


**Figure S3.** Detailed Analysis of CLO Responses to Key Therapeutic Agents Over Time.

(A) Temporal Response of CLOs to Ibrutinib Across Three PCNSL Patients from Day 2 to Day 20. Data on organoid size are presented as means ± SE, derived from three independent experiments.

(B) Time-Course Response of CLOs to Methotrexate in Three PCNSL Patients from Day 2 to Day 20. Organoid size metrics are shown as means ± SE, based on three independent trials.

(C) Longitudinal Response of CLOs to Dexamethasone Across Three PCNSL Patients from Day 2 to Day 20. Data on organoid dimensions are presented as means ± SE, collected from three separate experiments.

(D) Time-Dependent Response of CLOs to Rituximab in Three PCNSL Patients from Day 2 to Day 20. Organoid size data are shown as means ± SE, compiled from three independent experiments.

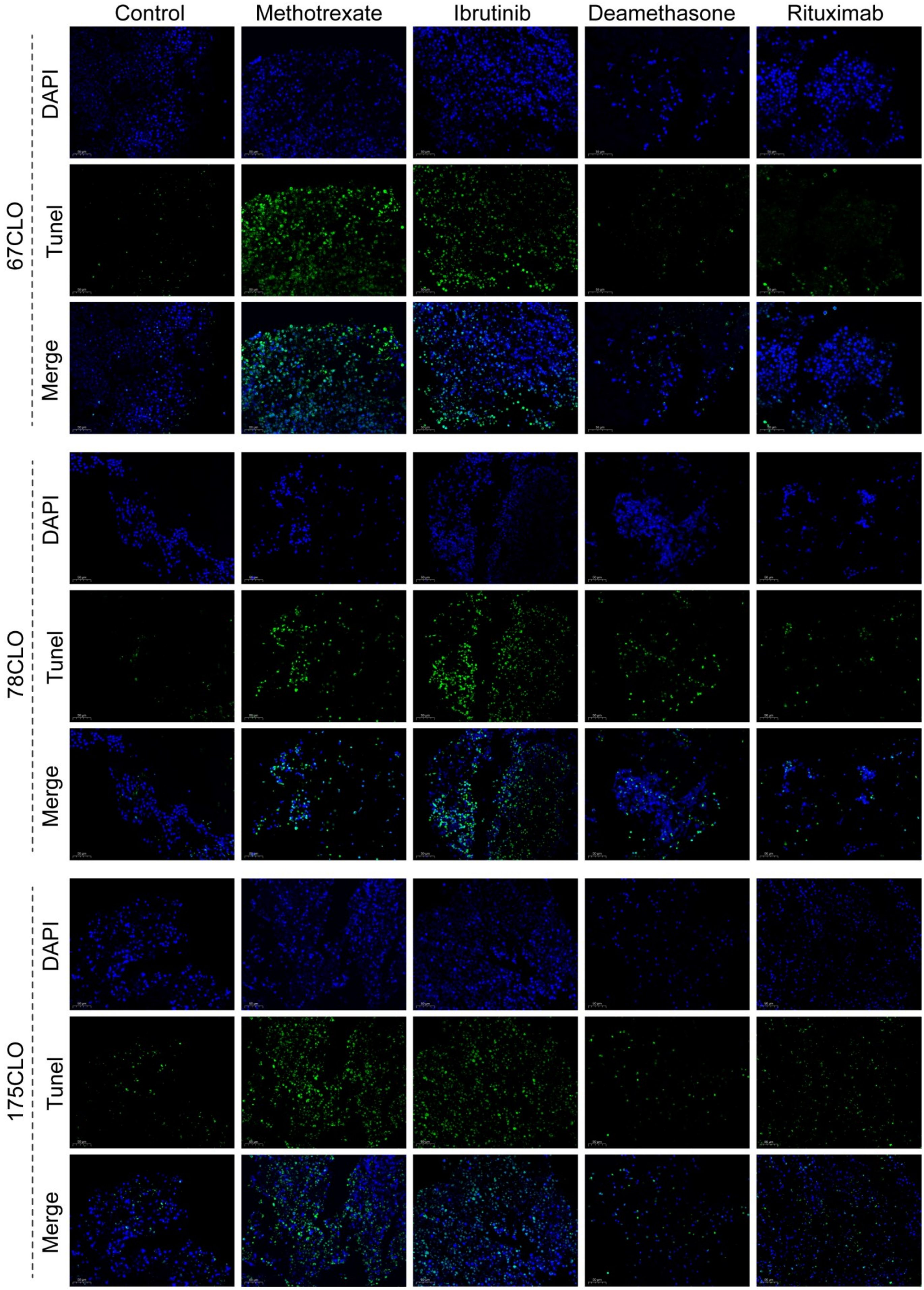


**Figure S4.** Assessing Drug Sensitivity (Ibrutinib, Methotrexate, Dexamethasone, and Rituximab) in PCNSL organoid (CLO) via TUNEL Assays.
